# Drug repurposing screens identifies compounds that inhibit α-synuclein oligomers’ membrane disruption and block antibody interactions

**DOI:** 10.1101/2022.10.06.511078

**Authors:** Arun Kumar Somavarapu, Giulia Kleijwegt, Madhu Nagaraj, Parvez Alam, Janni Nielsen, Daniel Otzen

**Affiliations:** Interdisciplinary Nanoscience Center (iNANO), Aarhus University, Gustav Wieds Vej 14, 8000 Aarhus C, Denmark

**Keywords:** Parkinson’s Disease, Vesicles, FDA, Small molecules, Inhibitors, Antibodies, Aggregation, neurons

## Abstract

Small soluble oligomers of the protein α-synuclein (αSO) have been linked to disruptions in neuronal homeostasis, contributing to the development of Parkinson’s Disease (PD). While this makes αSO an obvious drug target, the development of effective therapeutics against αSO are challenged by its low abundance and structural and morphological complexity. Here we employ two different approaches to neutralize toxic interactions made by αSOs with different cellular components. Firstly, we use available data to identify four neuronal proteins as likely candidates for αSO interactions, namely Cfl1, Uchl1, Sirt2 and SerRS. However, despite promising results when immobilized, all 4 proteins only bind weakly to αSO in solution in microfluidic assays, making them inappropriate for screening. In contrast, the formation of stable contacts formed between αSO and vesicles consisting of anionic lipids not only mimics a likely biological role of αSO but also provided a platform to screen two small molecule libraries for disruptors of these contacts. Of the 11 leads obtained in this way, 2 significantly impaired αSO contacts with other proteins in a sandwich ELISA assay using αSO-binding monoclonal antibodies and nanobodies. In addition, 5 of these leads suppressed α-synuclein amyloid formation. Thus a repurposing screening that directly targets a key culprit in PD pathogenesis shows therapeutic potential.

**Highlights:**

1. The toxic oligomer formed by α-synuclein (αSO) is an important drug target.
2. Neuronal proteins found by pull-down assays do not bind αSOs in solution.
3. Liposome assay identifies 7 approved drugs reducing αSO membrane disruption.
4. We identify different inhibitory mechanisms used by different compounds.
5. Two top drug hits disrupt αSO binding to oligomer-specific antibodies.

**Graphical Abstract:** 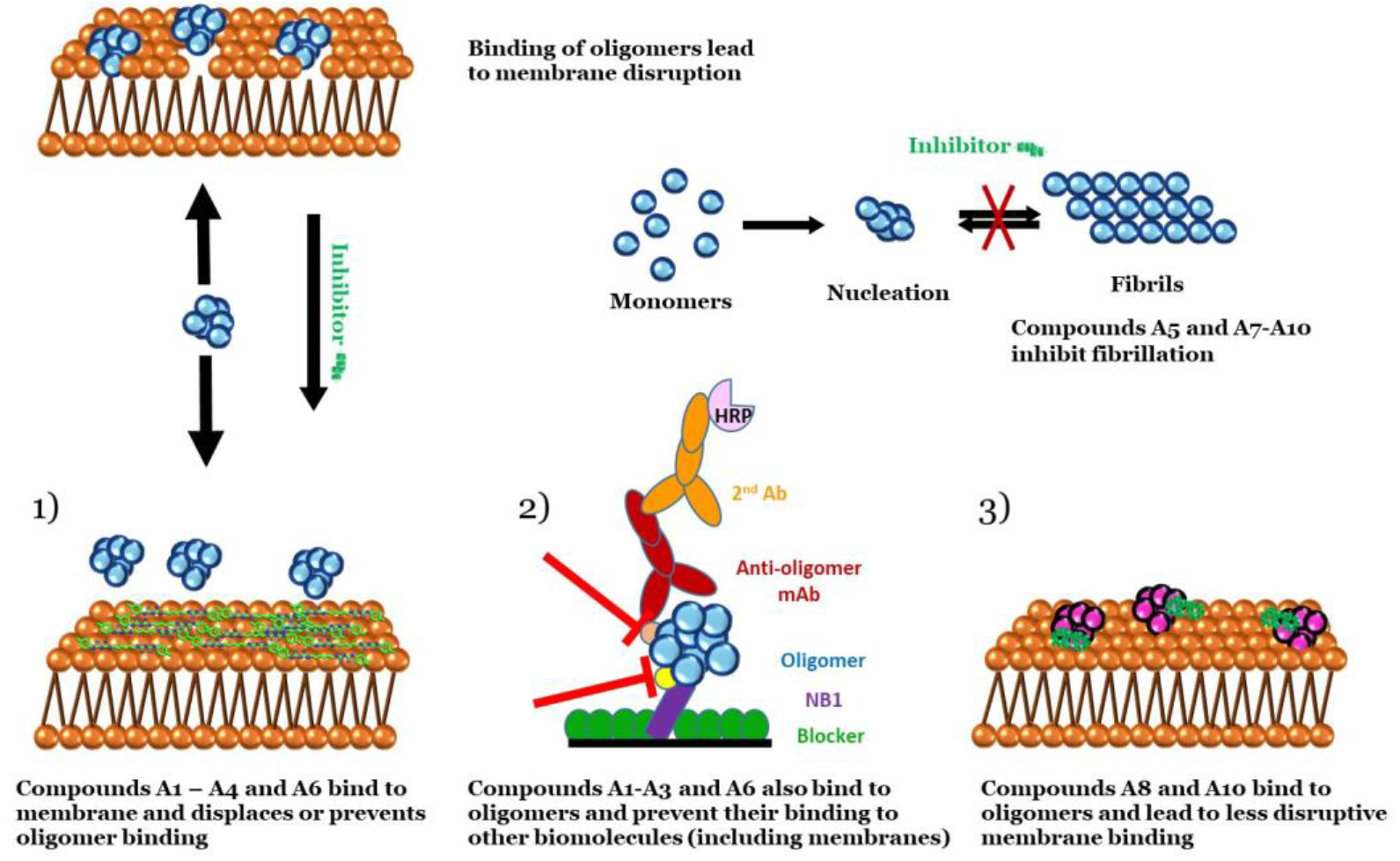

## INTRODUCTION

The protein α-synuclein (α-syn), widely expressed at presynaptic terminals of neurons, is associated with both sporadic and familial forms of Parkinson’s Disease (PD), the second most common neurodegenerative disorder^1^. α-syn is intrinsically disordered in the cytosol, and attains a helical conformation when bound to cellular membranes. Aggregation of α-syn to form small soluble cytotoxic oligomers (αSOs) and large fibrillary aggregates, manifested as intracellular inclusions called Lewy Bodies (LBs), are critical in the development of PD^2^ and are exacerbated by impaired autophagic and lysosomal clearance pathways ^3^. There is increasing evidence that αSOs are particularly toxic agents in this process^4^.

αSOs exhibit their toxicity through several intracellular mechanisms. They can interact with cell membranes and disrupt their structure, leading to disruption in cellular metabolism and homeostasis, increase in ROS production and ultimately neuronal death^5,6^. Recent solution and solid-state NMR studies identify two structural elements essential for membrane disruption, namely a highly exposed lipophilic N-terminal region of α-syn that promotes strong interactions to lipid membrane and a rigid β-sheet rich core that penetrates the lipid bilayer^7,8^. In addition, αSOs impair several cellular pathways. Thus binding of αSOs to the mitochondrial TOM20 receptor inhibits its interaction with the co-receptor TOM22, and impairs mitochondrial protein import, resulting in reduced respiration and increased reactive oxygen species^9^. Trapping of neuronal α3-Na^+^/K^+^-ATPase by both oligomeric and fibrillar α-syn clusters reduces the pumping efficiency and impairs Na^+^ gradient across plasma membrane^10^.

Recent electron microscopy and tomography studies of LBs have revealed α-syn-immunoreactive inclusions within the crowded environment of proteins, lipids, lysosomal structures and mitochondria^11^, highlighting that these aggregates are likely to have many interaction partners, both proteins and lipids. This is supported by proteomic screen approaches which identify intra-neuronal proteins that can bind preferentially to oligomer species^10,12,13^. These studies demonstrates the existence of α-syn conformation specific (monomer, oligomer or fibril) interactions, and further provide opportunities to study pathological processes specific for oligomers.

αSOs constitute a challenging drug target since they are only formed at low levels (*in vitro* typically 1-2% of total α-syn used^14-16^) and with a high variety of structural flexibility, ranging from a few well-defined core regions to completely disordered termini and with other segments showing intermediate dynamics^8,15,16^. This is not compatible with conventional structure-based drug design approaches. Nevertheless, numerous *in vitro* and *in vivo* studies have uncovered a variety of small molecule inhibitors of α-syn aggregation, including antibiotics, polyphenols, curcuminoids, quinones, aminosterol and dopamine analogs^17,18^. Besides aggregation inhibitors, natural compounds such as squalamine completely suppress the toxicity of αSOs in human neuroblastoma cells by inhibiting their interactions with lipid membranes^19^.

Current treatment of PD only targets dopamine related symptoms, and they do not improve or alter the progression of neuronal toxicity and cell death. We urgently need to identify compounds that protect or restore neuronal health^20^. Direct targeting of αSOs is a promising avenue for successful PD diagnostics and therapy. To develop a screening strategy which can target and disrupt complexes that involve αSOs, we need to understand the nature of these interactions and their sensitivity to disruption by *e*.*g*. small molecule compounds which can curtail αSO toxicity. Besides potential protein binding partners, αSO interactions with membrane are likely to be toxic to cell metabolism and homeostasis, making them important targets for small molecules that could ultimately have therapeutic effect.

We here used a double-pronged approach. Initially we focus on interactions of αSOs made with four neuronal proteins (Cfl1, Uchl1, Sirt2 and SerRS) which we identified as potential targets based on previous proteomics studies. Of these four proteins, Cfl1 is an essential regulator of cytoskeletal dynamics in cells. However, its inactivation by α-syn has pathological effects in PD. Neurons incubated with WT and A30P α-Syn presented a ∼2-fold increase in inactivate phosphorylated cofilin respectively compared to control^21,22^. Uchl1 is abundantly produced in brain and catalyzes the hydrolysis of ubiquitylated peptides. However, the autosomal dominant missense Ile93Met mutation, which causes a rare familial form of PD, decreases its ubiquitin hydrolase activity. On the other hand, it also exhibits ligase activity towards αSyn, leading to K63-linked polyubiquityl chains which lead to inefficient clearance of αSyn in PD models^23,24^. Sirt2 deacetylates histone and non-histone substrates, thus regulates a large spectrum of physiological processes. It has been reported to increase α-syn toxicity in PD models by deacetylation at K6 and K10 residues (which are endogenously acetylated in mouse brain), thus making it more prone to aggregation^25,26^. While SerRS catalyzes the attachment of serine to tRNA, its direct role in PD has not been explored. However, it has been reported to recruit Sirt2 to erase prior c-Myc-promoted histone acetylation^27^. However, we find that although all 4 protein ligands interact with αSOs when immobilized, they bind weakly to αSOs in solution. This makes them less useful as model systems to identify complex disruptors.

Therefore, our second strategy was to focus on αSOs’ ability to form stable complexes with DOPG liposomes. We used the αSO-DOPG system to screen two small-molecule collections of in total 2067 approved drug compounds from two different collections, and identified 7 compounds that reduce membrane disruption by oligomers. Besides inhibiting membrane binding, some of the hit compounds also completely inhibit amyloid fibril formation. Finally, we validate the utility of this approach by demonstrating that two of our top hits disrupt αSO binding to specific antibodies in a novel ELISA sandwich assay. These molecules establish the molecular basis for selective regulation of oligomer toxicity and could be the basis for therapeutic agents to suppress αSO-driven neurodegeneration.

## MATERIALS AND METHODS

### α-syn oligomer preparation

WT human α-syn was recombinantly expressed in *E. coli* Bl21 cells and purified by anion exchange chromatography as described ^16^. α-syn fractions were pooled and dialyzed against water, lyophilized and stored at -20°C. To prepare α-syn oligomers (αSOs), 10 mg of lyophilized α-syn was re-suspended into 1xPBS pH 7.4. The sample was then passed through 0.22 µm filters and incubated at 37°C for 4 h while shaking continuously at 900 r.p.m. After incubation, the solution was centrifuged at 14000 r.p.m for 5 min and the supernatant was loaded on to a Superdex 200 (10/300 GL) column. Oligomer fractions (eluting around 10-12 mL) were collected and stored at -20°C. Immediately before use, oligomer fractions were concentrated using Centricon filters with a 100 kDa cut-off. Oligomer concentration was determined by UV spectroscopy, using an ε_280_ of 0.412 mg^−1^ cm^-1^.

### Expression and purification of selected protein partners

Genes for the human proteins Cfl1 (residues M1-L166), Uchl1 (residues M1-A223), Sirt2 (residues M1-Q389), Sirt2_50 (residues S50-Q389) and SerRS (residues M1-A514) were prepared by Genscript (Piscataway, NJ), cloned separately into a pET30a(+) vector with the restriction sites *NheI* and *XhoI* and a C-terminal hexa-histidine tag. The Uniprot IDs of all four proteins are provided in Table 1. All genes were codon optimized for maximal expression in *E. coli* (GenScript). In each case, plasmids were transformed into the *E. coli* strain BL21 (DE3) cells separately for overexpression. Cells were grown in Luria-Bertani (LB) media and purified by nickel-affinity chromatography as described ^28-31^. Briefly, bacterial cells were grown in LB media containing 50 µg*/*ml kanamycin at 37°C with shaking to an OD_600_ of 0.6-1.0, expression was induced with 1 mM IPTG and the culture was incubated while shaking for 5 h. Cells were then harvested by centrifugation at 4000 rpm for 20 min and re-suspended in buffer A (20 mM Tris-HCl (pH 8.0) and 250 mM NaCl) with 10 mM imidazole, 0.1% v/v Triton X-100 and 1 tablet of EDTA-free protease inhibitor (Roche). The cell pellets from 4L of culture were lysed by sonication on ice and centrifuged at 12500 rpm for 40 min at 4°C. The resulting supernatants were loaded onto a Ni-NTA column and the column was washed with 10 column volumes of buffer A with 20 mM imidazole. The bound protein was eluted with buffer A with 250 mM imidazole. The fractions containing the protein were pooled, dialyzed into a buffer containing 10 mM HEPES (pH 7.4), 100 mM NaCl and 1 mM TCEP, and stored at -80°C.

**Table1.** Names and properties of the four proteins selected for αSO binding studies.

| Protein name | Gene name | Uniprot ID | # residues | Mass (KDa) | Function |
| --- | --- | --- | --- | --- | --- |
| **Cofilin-1** | Cfl1 | **P23528** | 166 | 18.5 | Binds to F-actin and regulates actin cytoskeleton dynamics |
| **Ubiquitin carboxyl-terminal hydrolase isozyme L1 (UCH-L1)** | Uchl1 | **P09936** | 223 | 24.8 | Processes ubiquitin precursors and ubiquitinated proteins |
| **NAD-dependent protein deacetylase sirtuin-2** | Sirt2 | **Q8IXJ6** | 389 (full-length), Sirt2_50 (50 - 389) lacking the nuclear export signal sequence | 43.2 (38.0 for Sirt2_50) | Deacetylates internal lysines on histones, alpha-tubulin as well as many key transcription factors |
| **Serine-tRNA ligase** | SerRS | **P49591** | 514 | 58.8 | Catalyzes the attachment of serine to tRNA<sup>Ser</sup> |

### Cell lysate preparation

SH-SY5Y human neuroblastoma cells (∼ 6 × 10^6^ cells) were washed with ice cold PBS twice. 2.5 mL of PBS was added to cell flask and cells were removed using a cell scraper. Cells were transferred to a falcon tube, centrifuged at 4000 rpm for 5 minutes, supernatant discarded and the cell pellet dissolved by suspension in 300 µL of N-PER™ Neuronal Protein Extraction Reagent (Thermo Scientific catalog number: 87792) lysis buffer. Subsequently, the suspension was gently mixed by inverting the tube up and down several times for 10 seconds and this process was repeated 3 times for 10 min. Cell debris was then pelleted by centrifuging the lysate for 10 min at 4 °C, 10,000 RCF in a tabletop centrifuge. The supernatant (which was used as cell lysate) was stored separately on ice and used within same day. Cells were kept on ice during the entire process. The lysate concentration was crudely estimated through absorbance at 280 nm (1 Abs ≡ 1 mg/mL).

### SPR analysis

Cfl1, Uchl1 and Sirt2 were immobilized on individual lanes of a CM5 sensor chip using amine coupling chemistry through immobilization solutions (EDC, NHS and EA from amine coupling kit). Running buffer contained 10 mM HEPES pH 7.4, 100 mM NaCl, 0.5 mM TCEP and 0.01% BSA. αSOs were injected at different dilutions over the active surfaces exposing the three binding proteins on individual lanes, and a blank surface control. Assays were performed at 25 °C on a Biacore 2000 instrument (GE Healthcare). Regeneration of the surface to remove bound analyte was carried out using 0.005% SDS. The sensorgram data was evaluated using the Biacore evaluation software (GE Healthcare). The kinetic data was fitted to a 1:1 binding model to give the equilibrium dissociation constant *K*_d_.

### Dot-blot binding assay

0.05–0.4 µg of each neuronal protein ligand was applied as 2 µL spots on three separate nitrocellulose membrane strips and allowed to dry. After drying, the strips were transferred to separate falcon tubes and non-specific sites blocked by soaking in 5 mL of 0.5% BSA for 1 h. The strips were then incubated overnight at 4°C with 100 μg of either α-syn monomers, oligomers or fibrils in 3 mL PBS solution. After incubation, the strips were washed three times with TBS-T washing buffer (0.05% Tween20 in TBS buffer), incubated with 3 mL of 1 µg/ml primary antibody (the monoclonal antibody 14-9E7-A1 obtained after immunization of mice with αSOs; this antibody binds strongly to αSOs but also recognizes monomers and fibrils (J. N. and D.E.O., unpublished data)) for 1 h at room temperature, washed three times with TBS-T and incubated with 20,000-fold diluted secondary antibody (goat anti-mouse coupled to horseradish peroxidase from Jackson Immunoresearch, Ely, UK) for 1 h at room temperature, washed and incubated with 2–3 mL TMB blotting solution for 5 min, washed with Milli-Q water and dried. The dried blots were imaged with a Geldoc Go imaging system (Biorad, Hercules, CA) and densitometrically analyzed with ImageJ^32^.

### Preparation of DOPG liposomes

To prepare pure liposomes of 100 nm diameter, 5 mg/ml of 1,2-dioleoyl-*sn*-3-phosphatidylglycerol (DOPG) was resuspended in 1xPBS buffer, subjected to 10 freeze-thaw cycles using liquid nitrogen and a 50 °C water bath and then passed through a mini-extruder 21 times using a 100 nm cut-off filter (Avanti Polar Lipids, Alabaster, AL). DOPG liposomes for calcein release experiments were prepared in the presence of 70 mM calcein (self-quenching concentrations). Calcein-filled liposomes were separated from free calcein using a PD-10 desalting column (GE Healthcare).

### Membrane permeabilization assay

Calcein release from calcein-filled vesicles upon membrane permeabilization was monitored by measuring the fluorescence at time 0 (F_0_) and after 1 h at 37°C (F) with excitation at 485 nm and emission at 520 nm for 1 h at 37 °C. Finally, Triton X-100 (0.1% (w/V)) was added to measure fluorescence corresponding to complete calcein release (F_max_). The % of calcein release is then determined as follows:

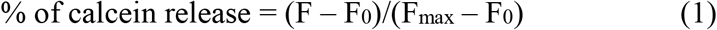

Dose-response calcein release for αSOs was quantified in presence of each of the hit compounds, and IC_50_ values were obtained by fitting dose-response curves to a sigmoidal model:

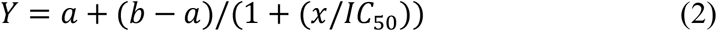

Here *x* is αSO concentration while *a* and *b* refer to bottom and top baseline levels of the curve.

### Small molecule libraries

Compound libraries were purchased from the Chemical Biology Consortium Sweden (CBCS) platform, which provides high quality bioactive chemical compounds. The chemical collection used in the screening contained 2067 unique pharmacologically active and chemically diverse compounds, derived from the Prestwick library (1200 FDA and EMA approved drugs) and the Biomol library (1031 compounds comprising neurotransmitter, nuclear receptor ligand, endocannobinoid and orphan ligand molecules). The total screening set thus contains 2231 compounds, of which 164 were duplicates. The compounds were provided at 10 mM stock concentrations in 100% dimethyl sulfoxide (DMSO), and were stored at -20 °C.

### Screening libraries using a calcein-based assay

For the primary compound screening, 150 nM αSO (concentration in α-syn monomer units) in PBS buffer was loaded on to 96-well plate (Nunc flat clear bottom, Thermo Fisher Scientific, Roskilde, Denmark), after which 5 μM of compounds diluted in 1xPBS and 0.05% DMSO was added. 5 µM each of EGCG and oleuropein were used as positive and negative control, respectively. The plates were sealed and incubated in a Clariostar fluorescence plate reader for 30 min at 37 °C and 2 s shaking every min. After incubation, F_0_ was recorded (λ_exc_ 485 nm; λ_em_ 520 nm) and 100 nm-size calcein-DOPG liposomes were added to each well at a final lipid concentration of 50 μM (monomer lipid units). Calcein release was measured for 1 hr at 37 °C with fluorescence recording after 2 s shaking every min (the average of the last 5 measurements was taken as F). Finally, 2 µl of Triton X-100 was added to each well to lyse vesicles, after which the fluorescence signal F_max_ was measured.

### Flow Induced Dispersion Analysis (FIDA)

Binding affinity studies of αSOs and DOPG liposomes were performed on a FIDA 1 instrument with a UV-LED fluorescence detector (Ex 480 nm/Em > 515 nm) (FidaBio ApS, Søborg, Denmark). Standard non-coated capillaries with inner diameter 75 µm, outer diameter 375 µm, total length 100 cm, length to detection window 84 cm were used for all experiments. The sample compartment and capillary housing is temperature controlled to 25 °C.

#### Alexa488 labeling of αSOs

To prepare indicator stock solution, αSOs was first concentrated to 40 µM (α-syn monomer units) using 100 kDa Centricon filters and phosphate-buffered saline (PBS) which is free of ammonium ions or primary amines, ensuring optimal labelling efficiency. The conjugation reaction was performed by incubating αSOs and Alexa-488 NHS Ester (Thermo-Fisher, Waltham, MA) at a 1:3 protein:dye molar ratio for 1 h at room temperature and then desalted on a PD-10 desalting column (GE Healthcare), where the free dye and the conjugated Alexa488-αSOs were separated. The NHS Ester group of Alexa-488 reacts with primary amines on proteins. The concentration was calculated according to the absorbance at 280 nm (αSOs) and 488 nm (Alexa-488). The degree of labelling was ∼0.6 labels per monomer protein, corresponding to ∼18 Alexa488 per αSO (for comparison, the recommended level of labelling for IgG molecules, a third of the size of an αSO, is 4-9 per IgG _33_).

#### Binding of αSOs to DOPG vesicles

1 µM αSOs-Alexa488 was prepared in assay buffer (PBS with 0.05% bovine serum albumin), diluted to a fixed indicator concentration of 150 nM (α-syn monomer units) and used with an analyte (DOPG liposome) concentration range of 0-150 µM. The apparent dissociation constant *K*_*d*_ was obtained by fitting the fraction of liposome bound αSOs (*Y*) to the concentration of DOPG lipids using a conventional binding isotherm that also includes a parameter (*slope*) to account for unspecific binding:

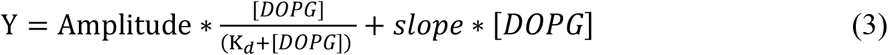

#### αSO-vesicle binding Inhibition assays

Stock solutions of 1 µM αSOs-Alexa488 and 50 µM compound were diluted to a fixed concentration of 150 nM and 5 µM, respectively, after which 500 µM DOPG liposomes was added to reach a fixed analyte concentration of 50 µM. All samples were preincubated for 15 min to attain equilibrium prior to analysis in the presence of DOPG liposomes and analyzed as follows: First, the capillary was rinsed and equilibrated with 1 M NaOH and assay buffer at 3500 mbar for 45 and 120 s, respectively. Subsequently, the analyte solution (DOPG liposomes) was injected into the capillary at 3500 mbar for 20 s followed by the indicator sample (αSOs-Alexa488 with/without compound, mixed with analyte solution) at 50 mbar for 10 s. Finally, the indicator sample was mobilized towards the fluorescence detector with analyte sample at 100 mbar for 600 s. All raw data taylorgrams were processed using FIDA data analysis software to calculate the hydrodynamic radius *R*_h_.

#### Effect of lead compounds on αSOs

To probe the lead compounds’ ability to aggregate or disaggregate αSOs, 150 nM αSO samples were pre-incubated with 5 µM lead compounds for 1 h at room temperature, after which analyte solutions with lead compounds alone were used to mobilize the indicator sample towards the detector.

#### Binding of αSOs to neuronal proteins

A stock solution of 1 µM αSOs-Alexa488 was prepared in working buffer and subsequently diluted to a fixed indicator concentration of 200 nM in the analyte concentration range of 0-150 µM protein ligands. All samples were pre-incubated for 15 min, to attain equilibrium prior to analysis, and the samples were analyzed as follows: First, the capillary was rinsed and equilibrated with 1 M NaOH and assay buffer, at 3500 mbar for 45 s and 120 s respectively. Subsequently 4 µL of analyte solution (protein ligands) at 3500 mbar for 20 s, followed by injection of 39 nL of αSOs-Alexa488 (pre-incubated with analyte) at 50 mbar for 10 s, which was then mobilized towards the detector with the analyte solution at 400 mbar for 180 s at 25 °C, pH 7.4.

### Ultracentrifugation for measuring compound – vesicle binding

To measure the membrane binding affinity of different compounds, 100 µM of each hit compound was incubated with and without 1 mM of DOPG vesicles for 30 min at room temperature. To separate the vesicle-bound compound (pellet) from free compound (supernatant), the samples were then centrifuged for 1 h at 40,000 rpm using Beckman Optima MAX-XP ultracentrifuge at 25°C. Subsequently, the absorption of compounds was measured where they displayed the highest absorption at 224 nm and 260 – 310 nm using Nanodrop. The % of compound bound to vesicles is then determined as follows:

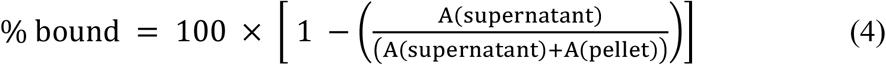

### Aggregation kinetics of α-syn in presence of inhibitors

A stock solution of α-syn was prepared by dissolving freeze-dried α-syn in PBS buffer (pH 7.4), after which it was filtered through a 0.22 µm nylon filter. Stock solutions of lead compounds in DMSO were prepared at 20 mM. Aggregation kinetics assay conditions include 20 µM α-syn, 20 µM compound, 0.1 % DMSO, 40 µM ThT, pH 7.4. 20 µM EGCG at similar assay conditions was used as positive control. The samples were loaded into a 96-well Nunc plate with 150 µl assay solution in each well. Plates were sealed with clear sealing tape. The change in the ThT fluorescence signal with time was monitored using a Clariostar plate reader under shaking conditions at 300 rpm (10 min shaking in 12 min measurement intervals) and 37 °C. ThT signal was measured with λ_exc_ 448 nm and λ_em_ 485 nm. All runs were recorded in duplicates and the average run is shown with error bars.

### Sandwich ELISA assay using specific antibodies and nanobodies

A High Bind ELISA plate (Sarstedt, Nümbrecht, Germany) was coated with 60 μL of 5 μg/mL anti-αSO nanobody (produced by standard llama immunization with αSOs followed by cloning of B-cells and selection by bacteriophage display^34^ (J.N. and D.E.O., unpublished results), incubated overnight at 4℃, emptied by inversion and gentle tapping on a table and blocked with 75 μL 2% BSA in 1x PBS for 30 min at 37℃. Then the plate was washed on a Intelispeed Washer IW-8 (BioSan, Riga, Latvia) with 0.05% Tween20 in 1x PBS three times and incubated with 50 μL 2 μg/mL αSO for 1 hour at 37℃. The plate was again washed, and then 25 μL compound was added (10 mM EGCG (Sigma-Aldrich, ≥95%), 1 mM A1 to A5 (Green Pharma), 1 mM A6-A10 (Merck) for 30 min at 37℃ (see Table 2 for details on the compounds). Then 25 μL of 1 μg/mL primary monoclonal anti-αSO antibody (14-9E7-A1) was added and the plate was incubated for 1 hour at 37℃, washed three times, and 50 μL secondary Jackson GAM-HRP antibody (1:20000) was added for 1 h at 37℃. The plate was again washed, and 50 μL TMB One (3,3’,5,5’-tetramethylbenzidine) was added for 30 min at 37℃ while keeping in the dark. The reaction was stopped by adding 100 μL 0.5 M sulfuric acid. The absorption was measured at 450 and 620 nm on a HiPo Microplate Photometer MPP-96 (BioSan, Riga, Latvia).

**Table 2.** The 11 best compounds from primary compound screening using calcein assay.

| Compounds | Sub-library | Compound trivial name | Normalized values<sup>a</sup> | Pharmaceutical activity |
| --- | --- | --- | --- | --- |
| A1 | Prestwick | Chlorhexidine | 0.03 | Antimicrobial and antiseptic |
| A2 | Prestwick | Methyl benzethonium | 0.03 | Surfactant, antiseptic, and antimicrobial |
| A3 | Prestwick | Benzethonium | 0.03 | Surfactant, antiseptic, and antimicrobial |
| A4 | Prestwick | Pinaverium | 0.14 | Spasmolytic agent used for gastrointestinal disorders |
| A5 | Prestwick | Metacycline | 0.02 | A tetracycline antibiotic, antimicrobial action |
| A6 | Biomol | Methoctramine | 0.26 | Antagonize muscarinic receptors, studied to treat bradycardia |
| A7 | Biomol | Shikonin | 0.24 | Derived from the roots of the shikonin plant, antibacterial, anti-inflammatory and anti-tumor |
| A8 | Biomol | Cisplatin | 0.04 | Chemotherapy medication, binds to DNA and inhibits replication |
| A9 | Biomol | Carboplatin | 0.15 | Chemotherapy medication, binds to DNA and inhibits replication |
| A10 | Biomol | Adrenochrome | 0.09 | Produced by the oxidation of adrenaline, The derivative carbazochrome, is a antihemorrhagic |
| A11 | Biomol | Tyrphostin 47 | 0.14 | A protein tyrosine kinase inhibitor particularly potent against EGFR kinase activity |

Notes:
<sup>a</sup> Relative extent of calcein release. 0 indicates complete suppression of calcein release (0% calcein release) while 1 indicates no effect of the compound compared to αSO alone (which leads to 50% calcein release). Thus the actual extent of calcein release is the normalized value x 50%.

## RESULTS

### Filtering proteins from proteomics studies to identify suitable binding partners for αSOs

We start by selecting potential binding partners for αSOs based on three proteomics-based studies^10,12,13^. In these studies, purified αSOs on agarose beads were used as bait protein to capture potential binding partners in a neuronal lysate, after which the beads were spun down and the binding partners identified by mass-spectrometry. The collection of 138 proteins obtained from these independent studies is a useful starting point although no direct information about binding affinity is provided. To select the most appropriate binding partners and to minimize the practical challenges in purifying them, we limited ourselves to proteins < 500 residues which were not membrane-bound or parts of stable multi subunit complexes. This reduced the data set to 31 proteins. These proteins were then matched against Parkinson’s UK annotation datasets to filter disease relevant genes^35^ (**Fig. 1a**). The annotation dataset contains 48 high priority proteins encoded by familial PD genes or genome wide association studies, and 330 proteins from an extended list of proteins that interact with the high-priority proteins or play a role in PD-related biological processes. Apart from α-syn itself, none of the 138 proteins from the three proteomic-based studies were matched with the proteins in the high priority set. On the other hand, we found three matches in the extended Parkinson’s annotation list. Of these three proteins, we discarded Non-POU domain-containing octamer-binding protein (NONO)) due to a high amount (∼30%) of predicted disordered sequence and its reported tendency to aggregate rapidly after purification^36^, leaving Cofilin-1 (CFL1)^13^ and Ubiquitin carboxyl-terminal hydrolase isozyme L1 (UCHL1)^10^ for further studies. We could not identify any ligand from the third study^12^ that matches Parkinson’s annotation list after the first filtering step. However, we chose two proteins from this study, namely the NAD-dependent protein deacetylase sirtuin-2 (SIRT2) and seryl-tRNA synthetase (SerRS) based on their high preferential binding to oligomers compared to monomers^12^ (gauged as the ratio between the MS peak areas of peptides occurring in immunoprecipitation for monomers and oligomers). The four proteins chosen vary in size (18.5 to 58.8 KDa) (**Fig. 1b**) and function as summarized in **Table 1**. Initial expression attempts with full-length 389-residue Sirt2 led to low protein yields, so we made a new construct called Sirt2_50 (residues 50-389) where removal of both the N-terminal disordered and nuclear export signal peptide enabled production of higher amounts of protein^37^. In summary, all 4 binding proteins that are studied here have been implicated in neurodegenerative diseases and in contacts with αSOs, but their specific mode of interaction has not been studied, motivating further investigations.

**Fig. 1.**
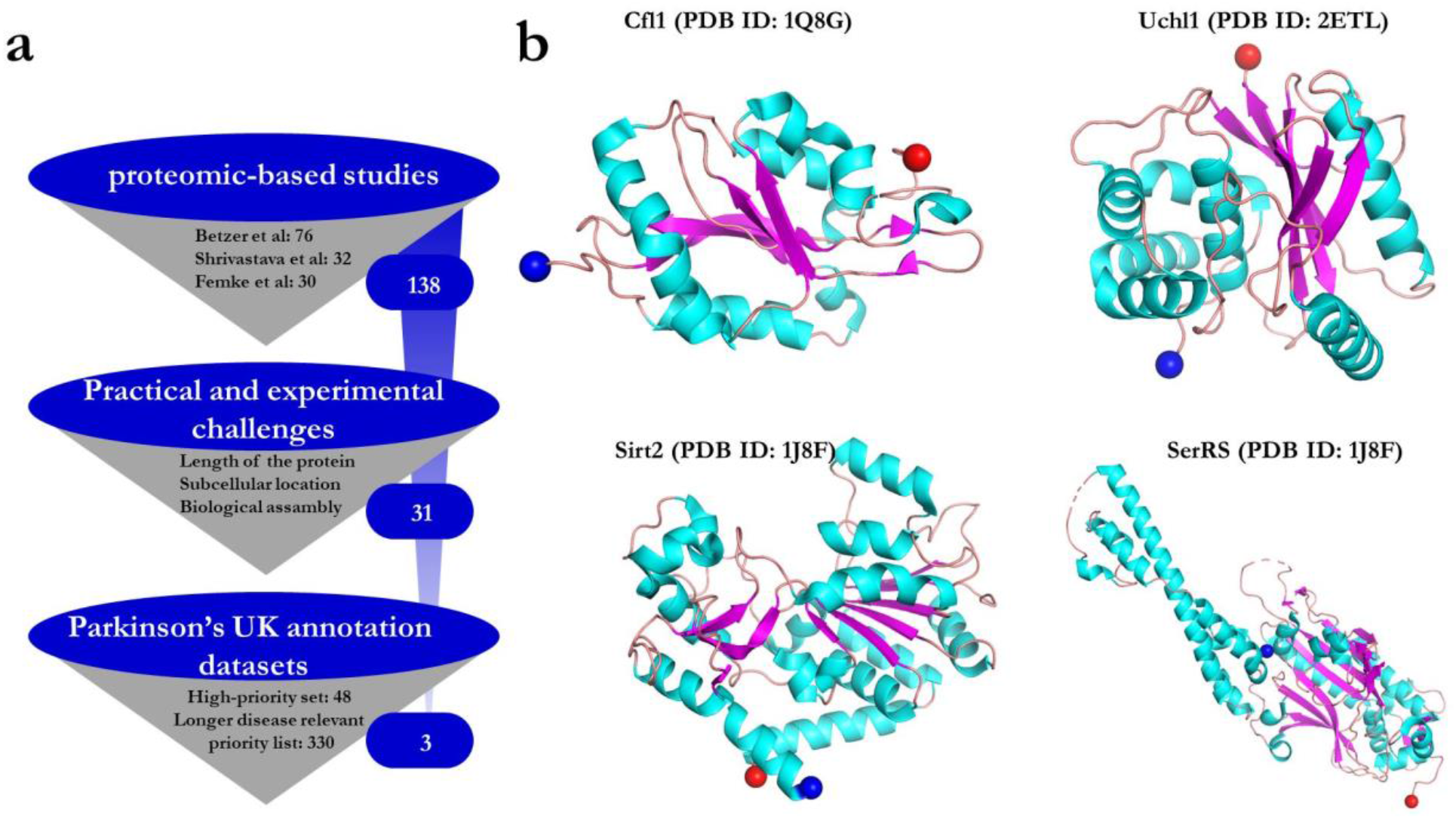
a.) Filtering steps used to select potential protein partners for interaction studies with αSOs. b.) PDB structures of all four protein ligands, colored according to secondary structure. The blue and red spheres represents N- and C-terminal ends of each structure, respectively.

### Immobilized and in-solution assays provide conflicting information about interactions of αSOs with four neuronal proteins

To investigate the binding specificity of the four identified protein ligands for αSOs, we first performed cross-reactivity studies using a dot-blot binding assay. We immobilized increasing amounts (0.05 to 0.4 µg) of ligands on nitrocellulose membranes and then added different species of α-syn (monomer, oligomer and fibril) in the mobile phase. Recombinant human Hsp90, which is known to interact with αSOs^38^, was used as a positive control. **Fig. 2a** shows the concentration dependent binding of neuronal proteins to different species of α-syn; the darker the dot, the stronger the binding. Using densitometric analysis, we calculated the degree of binding to different species of α-syn, based on the total intensity for all three species combined for each individual ligand (**Fig. 2b**). Each dot intensity is represented as the percentage of all the measured intensities (monomer + oligomer + fibril) for a particular ligand at that ligand concentration. HSp90, the positive control, showed the expected preferential binding to αSOs. Of the 4 ligands, only Cfl1 and SerRS showed higher binding to αSOs than to α-syn monomers and fibrils. Uchl1 bound most strongly to fibrils while both Sir2 and and Sirt2_50 bound to a similar extent to all three species. To confirm this binding profile, we turned to surface plasmon resonance (SPR). We chose Cfl1, Uchl1 and Sirt2 due to their different αSO binding profiles in the dot-blot assay. The 3 proteins were immobilized separately on individual lanes of a CM5 chip, after which we measured the binding kinetics of αSOs at various αSO concentrations (**Fig. 2c**). All proteins showed a concentration dependent increase in binding signal (though the signal was weak for Uch1), leading to estimated affinity constants of 0.29 µM, 0.15 µM and 0.6 µM for Cfl1, Sirt2 and Uchl1, respectively.

**Fig. 2.**
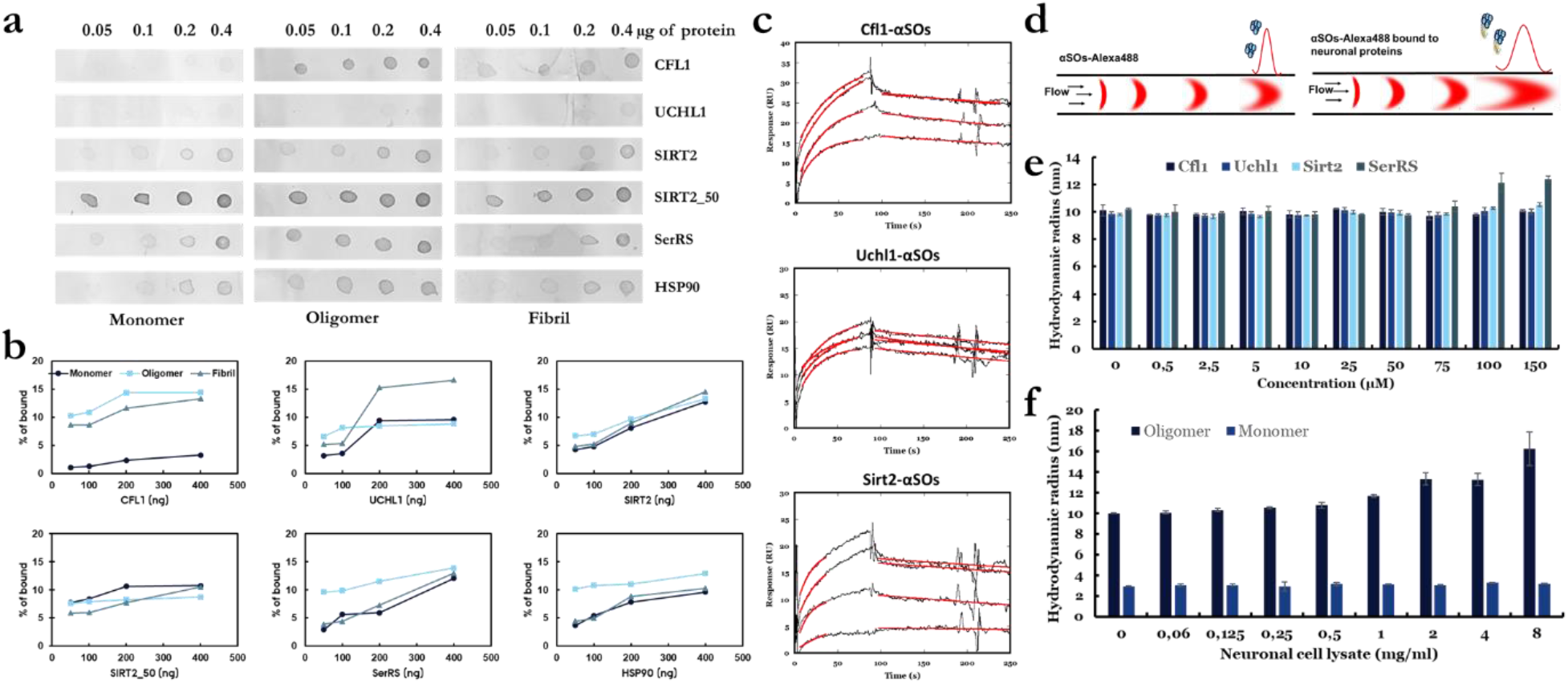
Characterizing the interactions of αSOs with neuronal proteins using various biophysical techniques. a) Dot blot binding assay using 0.05-0.4 µg of immobilized neuronal proteins exposed to 33 µg/ml α-syn in the soluble phase. b) Binding propensities of each ligand to different species of α-syn calculated using densitometric analysis (ImageJ software. Values are shown as percentages of the total intensity of all 12 binding dots for each ligand in panel a. c) Binding kinetics of αSOs measured by SPR analysis. 1-15 µM of αSOs were passed over sensor surfaces immobilized with Cfl1, Uchl1 and Sirt2 on separate lanes. Sensorgrams show the time curves of the SPR binding signal where the best fits to the binding region using a 1:1 binding model are provided in red. d) Principle of the use of FIDA to characterize αSOs – ligand interactions. e) *R*_h_ of αSOs-Alexa488 after binding to protein ligands as a function of ligand concentration. f) *R*_h_ of αSOs-Alexa488 and αSMs-Alexa488 after binding to different concentrations of neuronal cell lysate.

Immobilization of proteins potentially complicates interpretation of binding data. Therefore, to determine the binding affinity of these proteins to αSOs in solution, we turned to Flow-Induced Dispersion Analysis (FIDA), a microfluidic technique in which the hydrodynamic radius R_h_ of a labelled biomolecule (the indicator) can be obtained from its diffusion behavior (Taylor Dispersion Analysis) in a thin capillary ^39^ (**Fig. 2d**). Binding to other biomolecules of significant size will increase apparent indicator R_h_. The indicator R_h_ is measured in a titration series with varying concentrations of unlabeled binding partner (analyte) and the resulting binding curve is used to determine the binding affinity (*K*_d_) and complex size.

We used Alexa488-labelled αSOs as indicator in combination with unlabeled ligands. Pre-incubation mixing mode of samples were used where αSOs and a given ligand were pre-incubated for 15 mins prior to the measurements, and subsequently analyzed. In practice the solution was injected onto the capillary as a plug surrounded by buffer containing the corresponding concentrations of ligand sample, allowing the maintenance of a constant concentration of ligands, which is required for proper equilibrium measurements. The R_h_ of αSOs alone was 10 ± 0.2 nm, which is similar to the R_h_ of ∼11 nm that we previously reported based on SEC-MALS and SAXS^16^ and confirms the structural integrity of the αSO. Using FIDAbio’s PDB R_h_ predictor tool, we predicted the R_h_ of the four ligands, which were 2.3 nm (Cfl1), 2.4 nm (Uchl1), 2.8 nm (Sirt2) and 3.6 nm (SerRS). The R_h_ of αSO is expected to increase to 12 – 13 nm upon binding to ligands, and this increase is is well within the limits of detection (since increases as low as 0.5 nm should be measurable). However, we saw little change in R_h_ when titrated with up to 150 µM Cfl1, Uchl1 and Sirt2 (**Fig. 2e**). Only for SerRS was there a small but significant increase in size of oligomer (from 10 to ∼12.5 nm) and this only took place at high (100-150 µM) concentrations. This observation suggests that the binding affinity of ligands to αSO in solution is very weak and well below that indicated by dot-blot assays and measured by SPR.

### FIDA reveals cross reactivity of αSOs with functional neuronal proteins in cell lysates

While the protein ligands studied here did not show strong binding to αSOs in solution, we investigated whether there might be other neuronal components that could bind to αSOs in solution under physiological conditions, *i*.*e*. in a cell-like environment crowded with neuronal proteins. Different concentrations (0.06-8 mg/ml) of freshly prepared human neuroblastoma cell lysate were incubated with Alexa488 labelled αSOs and Alexa488 labelled α-syn monomer (αSMs-Alexa488) and the R_h_ values determined by FIDA. Remarkably, while there is no change in the size of monomer, significant increase in the average size of oligomer (10 nm to 16.2 nm) was observed (**Fig. 2f**). While FIDA does not allow us to identify the binding partner(s), our data clearly show that there are in-solution interaction partners for αSOs in the lysate, consistent with the ability to identify interaction partners in proteomic studies^10,12,13^.

### Protein ligands either inhibit or stimulate α-syn aggregation

Despite the weak interactions of αSOs with the 4 binding partners in solution and the caveats associated with their identification by immobilization techniques, their ability to interact with immobilized αSOs implies a potential to affect α-syn aggregation. We therefore followed aggregation kinetics of 20 µM α-syn monomer in presence of different concentrations of the four ligands (Cfl1, Uchl1, Sirt2_50 and SerRS) by ThT fluorescence (**Fig. 3**). On its own, α-syn alone at 20 µM concentration showed an aggregation half time of 44 h. As little as 1 µM Cfl1 led to a dramatic reduction in aggregation. The effect reached saturation at sub-stoichiometric concentrations of 10 µM. In contrast, the other three ligands (Uchl1, Sirt2_50 and SerRS) promoted aggregation of α-syn. Thus 5 µM Sirt2_50 and Uchl1 SerRS reduced half-times of aggregation 2 and 3-fold respectively, while as little as 1 µM of SerRS ligand reduced half time of α-syn fibrillation more than 3-fold. This indicates a strong impact of the binding partners on α-syn fibrillation which is not related to αSO formation, given that αSOs are off-pathway to fibrillation^16^.

**Fig. 3.**
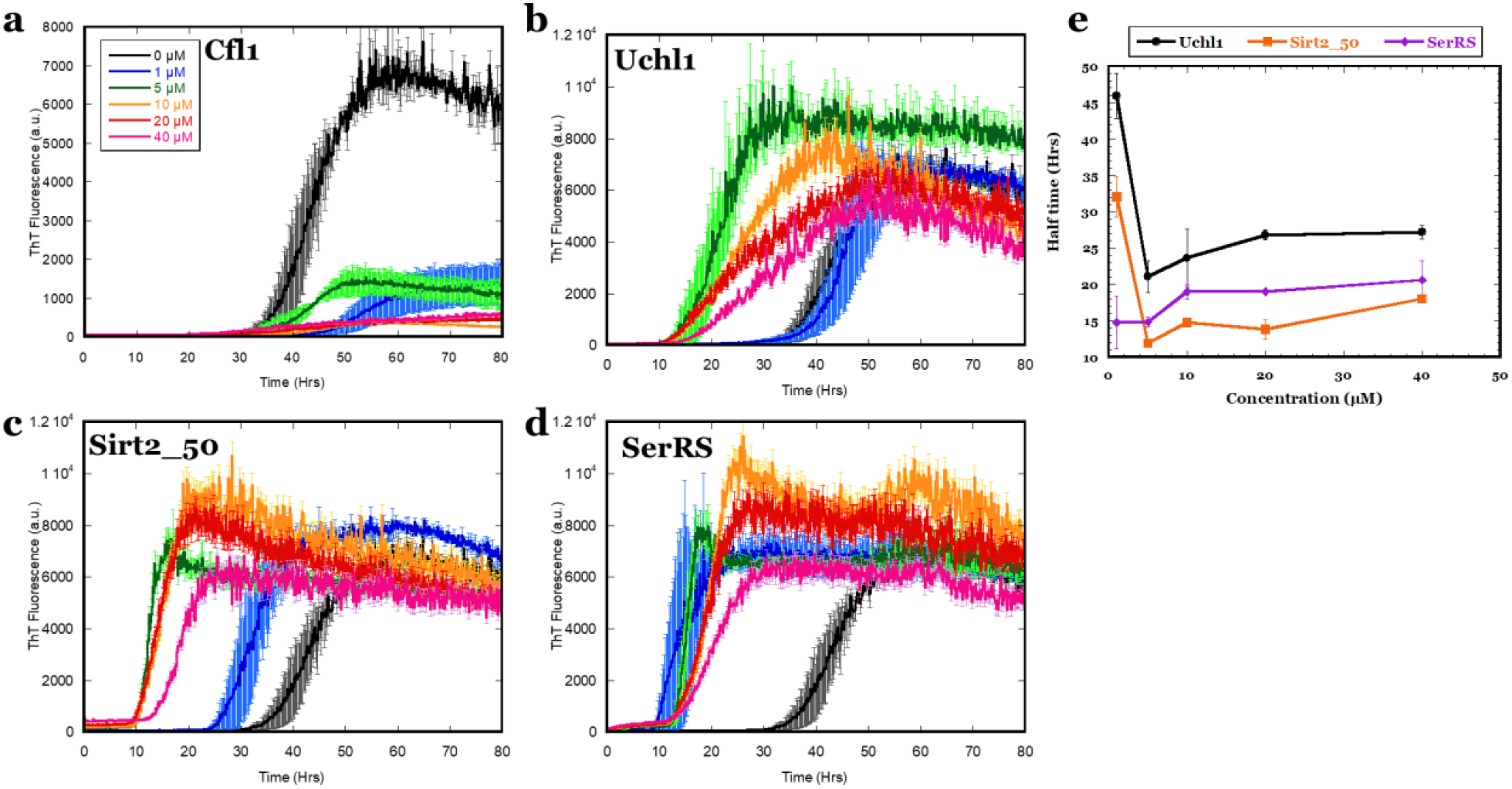
Tht aggregation kinetics of α-syn in the presence of four different protein ligands a.) Cfl1, b.) Uchl1, c.) Sirt2_50 and d.) SerRS. e.) t_1/2_ of aggregation as a function of ligand concentration for the 3 αSO ligands which show variation in lag times rather than ThT end-point levels. Colour codes in panels b-d as indicated in panel a.

### Using FIDA to characterize oligomer-membrane interactions

Given the weak interactions of the proposed binding partners to αSOs, we instead turned to the αSO-membrane system as an approach to identify small molecules that disrupt αSO interactions. To elucidate the cytotoxic interactions of αSOs bound to cell membranes, we prepared membrane-mimicking liposomes made of DOPG lipids and titrated them against αSOs using FIDA. As for the protein ligand studies, αSOs-Alexa488 and DOPG liposomes were co-incubated for 15 min prior to the measurements, and subsequently analyzed with corresponding concentrations of liposome sample in the surrounding solvent (**Fig. 4a**).

**Fig. 4.**
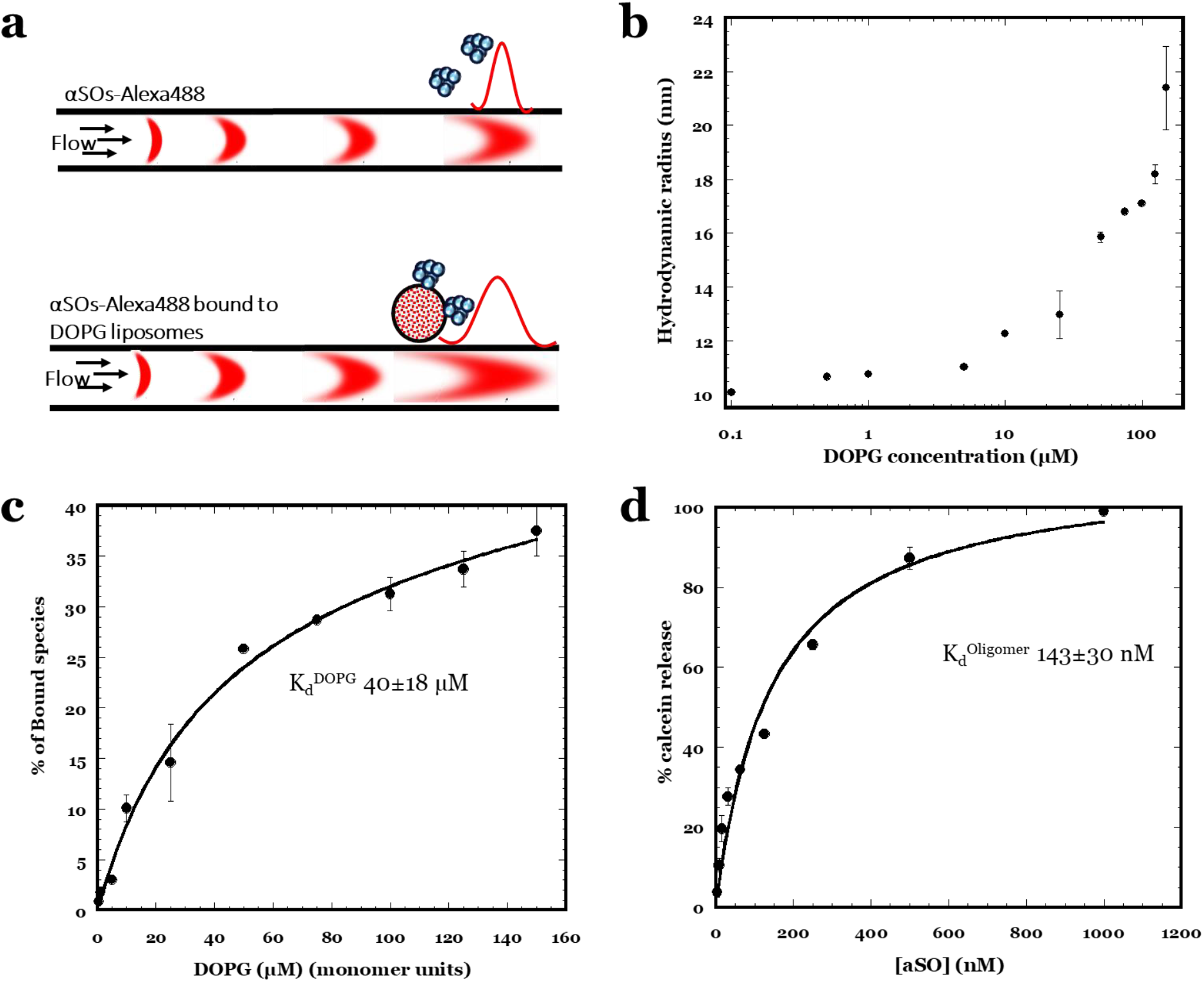
a.) Principle of the use of FIDA to characterize αSOs – DOPG liposome interactions. b.) *R*_h_ of αSOs-Alexa488 after binding to DOPG liposomes as a function of DOPG lipid concentration. c.) The apparent dissociation constant K_d_ (where lipid concentration is expressed in monomer units) was obtained using data from panel b which were transformed and fitted with eq. 3 (including a term for unspecific binding) as described. d.) Extent of release of calcein from DOPG vesicles incubated with different concentrations of αSO. Data fitted to a binding isotherm with an amplitude of 100.

*R*_h_ of αSOs-Alexa488 in the absence of liposomes is 10.15 ± 0.02 nm (consistent with our previous measurements), while that of the α-syn-Alexa488 monomer is 3.0 ± 0.03 nm, in good agreement with previous single-molecule fluorescence correlation spectroscopy (FCS) measurements^40^. DOPG liposomes extruded to an expected R_h_ of 50 nm were then titrated against αSOs-Alexa488. As the concentration of liposomes is increased, the apparent size of αSOs-Alexa488 increases due to binding. The apparent R_h_ of liposome bound αSOs-Alexa488 were determined using three-species taylorgram fitting where the size of the one unknown species (here αSOs) is determined by fixing the sizes of two known species in the sample, *i*.*e*. free label (0.6 nm) and monomers (3.0 nm). Fitting the FIDA data thus provides the average αSO size, *i*.*e*. a weighted average of free and liposome-bound αSOs (**Fig. 4b**). Furthermore, to differentiate the oligomer species into liposome bound and non-bound, we fixed the size of pre-determined αSOs as 10.2 nm and liposome bound αSO as 60 nm (assuming 1:1 binding stoichiometry) and fitted the FIDA data again. The resulting plot of the percentage bound species as a function of lipid concentration (**Fig. 4c**) shows a steep rise at low lipid concentrations followed by a more shallow increase at higher. When fitted with a model that includes specific binding (a hyperbolic relationship) along with a weaker and more unspecific binding (linear relationship), we obtain an apparent lipid binding affinity 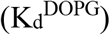 of around 40 µM (**Fig. 4c**). However, the actual affinity of oligomers for vesicles is likely higher (see below and **Fig. 4d**).

### Screening small molecule libraries to inhibit oligomer-membrane interactionsxs

To identify compounds that inhibit αSO permeabilization of membranes, we first optimized the calcein assay conditions (illustrated in **Fig. 5a**) for screening. When we add increasing amounts of αSO (3.9-1000 nM) to a constant concentration of DOPG liposomes (50 µM), there is a gradual increase in calcein signal with ∼50 % calcein release around 150 nM αSO (**Fig. 4d**). The data can be fitted to a binding curve with an amplitude of 100% to give an apparent 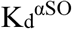 of 143 nM. We therefore decided to use 150 nM αSO and 50 µM DOPG as standard conditions under which to screen two datasets of compounds that covers both FDA approved (Prestwick library) and clinical-stage biologically active drugs (Biomol library). We used EGCG and Oleuropein as positive and negative controls, given that EGCG completely inhibits calcein release^41^, whereas Oleuropein displays weak or no inhibitory effect^42^.

**Fig. 5.**
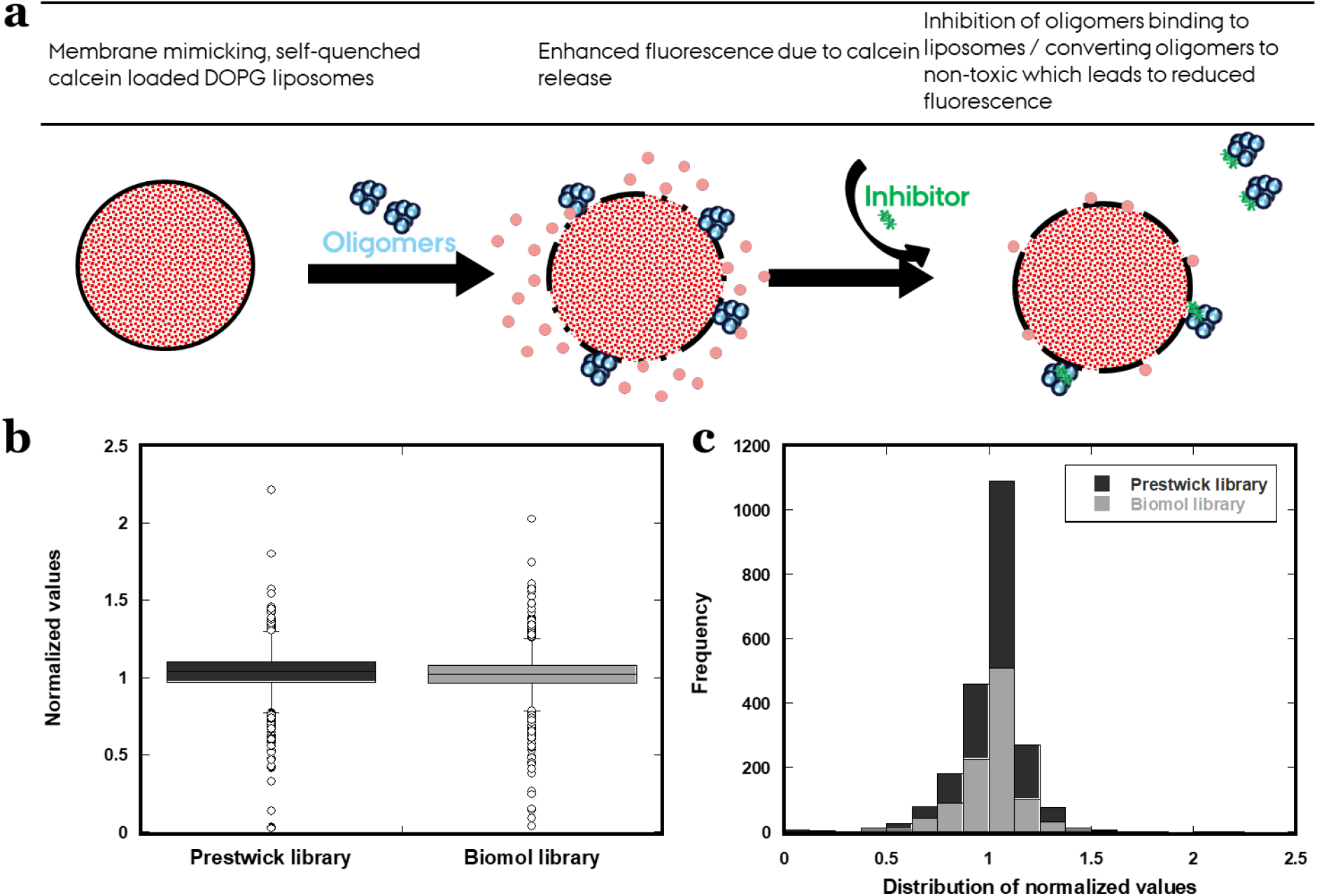
a.) Illustration of the membrane permeabilization assay using calcein-filled DOPG liposomes, b.) Boxplot and c.) Histogram showing the distribution of normalized calcein release values form primary compound screening. See text for definition of normalized values.

Calcein release values of all the compounds were normalized according to control (αSOs alone without compounds) (**Fig. 5b and 5c**) i.e. ∼50% of calcein release. Here 0 refers to complete suppression of calcein release (that is, 0% calcein release), 1 refers to no effect of the compound compared to αSO alone (that is, 50% calcein release) and >1 refers to an increase in calcein release caused by the compound (>50% calcein release). Out of 2067 unique compounds studied, 24 compounds inhibited the calcein release by at least 50% (that is, they led to 25% calcein release or less) compared to the absence of compounds. To identify the most potent compounds for further studies, we set the threshold at 15% calcein release (i.e. a reduction by 35% or more compared to the absence of compounds), resulting in 11 (0.5%) hits (**Table 2, Fig. 6**). Of these, five were from the Prestwick library and six from Biomol. While most of the remaining compounds did not significantly affect the calcein release, ∼12 compounds (those with normalized values of > 1.5) increased the release of calcein by more than 50% on top of the existing 50% by αSOs alone i.e. at least 75% calcein release.

**Fig. 6.**
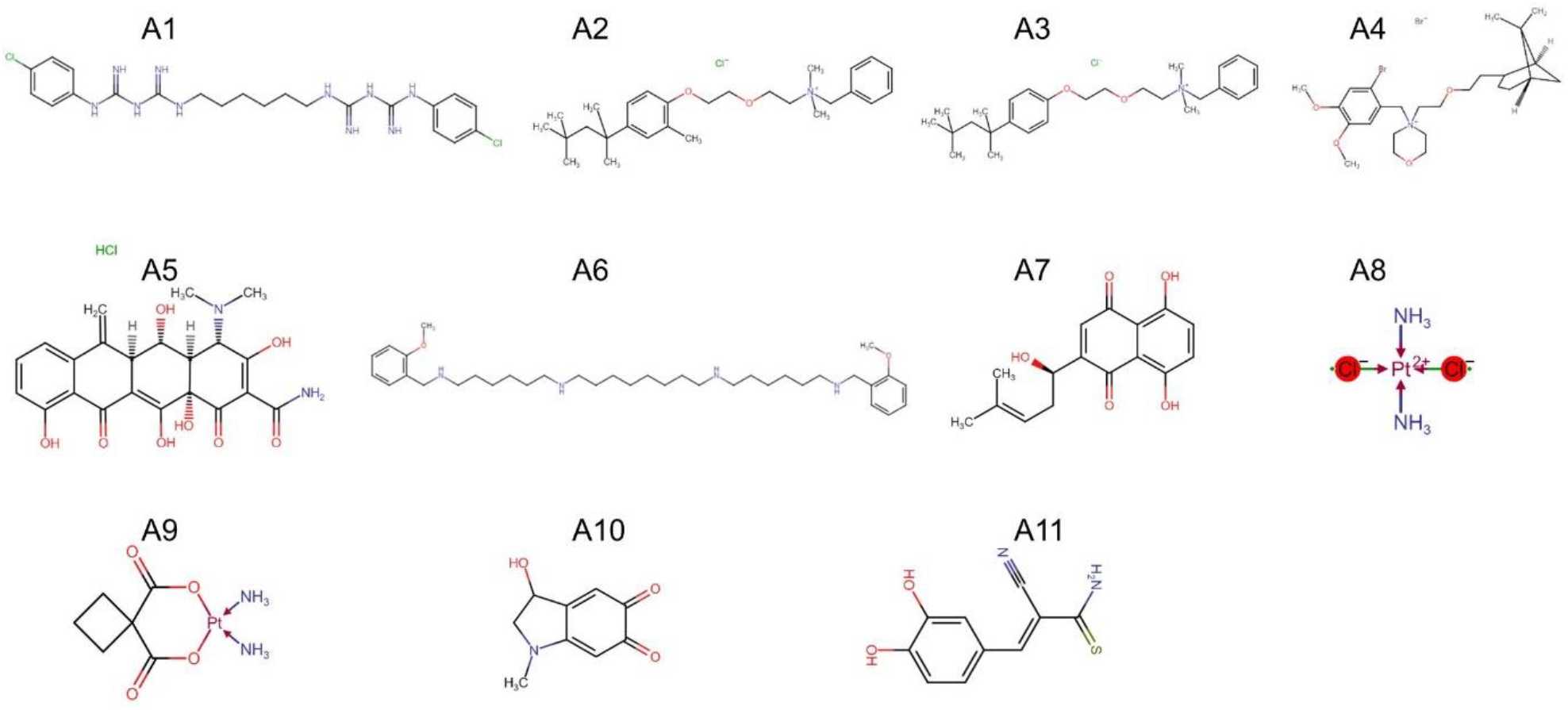
The 11 compounds which reduce αSO-driven membrane permeabilization to at most 15% calcein release. Details provided in **Table 2**.

### Dose-response curves rank the different hit compounds

We constructed dose-response curves for 10 of the 11 best compounds with our calcein assay, (compound A11 was not available for purchase from vendors). Six compounds (A1 – A4, A6 and A8) maintained the inhibition of vesicle permeabilization as expected from primary compound screening. Of the remaining 4, compound A10 displayed lower efficacy at concentrations used for primary screening (5 µM), while compounds A5, A7 and A9 failed to show any response even at 20 µM, indicating that they were false positives in terms of membrane interactions. For the six compounds which showed an effect, we obtained IC_50_ values by fitting data to eq. 2 (**Fig. 7a and 7b**). Compounds A1 – A4 displayed similar IC_50_ values (1.65 – 2.65 µM); the highest inhibitory effect was shown by A6 and A8 with IC_50_ values of 0.32 µM and 1.05 µM respectively. Both EGCG and PGG plant polyphenols that were included in the study for comparative analysis exhibited a much more pronounced inhibitory effect than the hit compounds, giving IC_50_ values of 0.08 µM and 0.01 µM respectively. Overall, the hit compounds inhibits membrane permeabilization of αSOs in the ranking order of A6 > A8 > A1 – A4 > A10 (**Fig. 7c**).

**Fig. 7.**
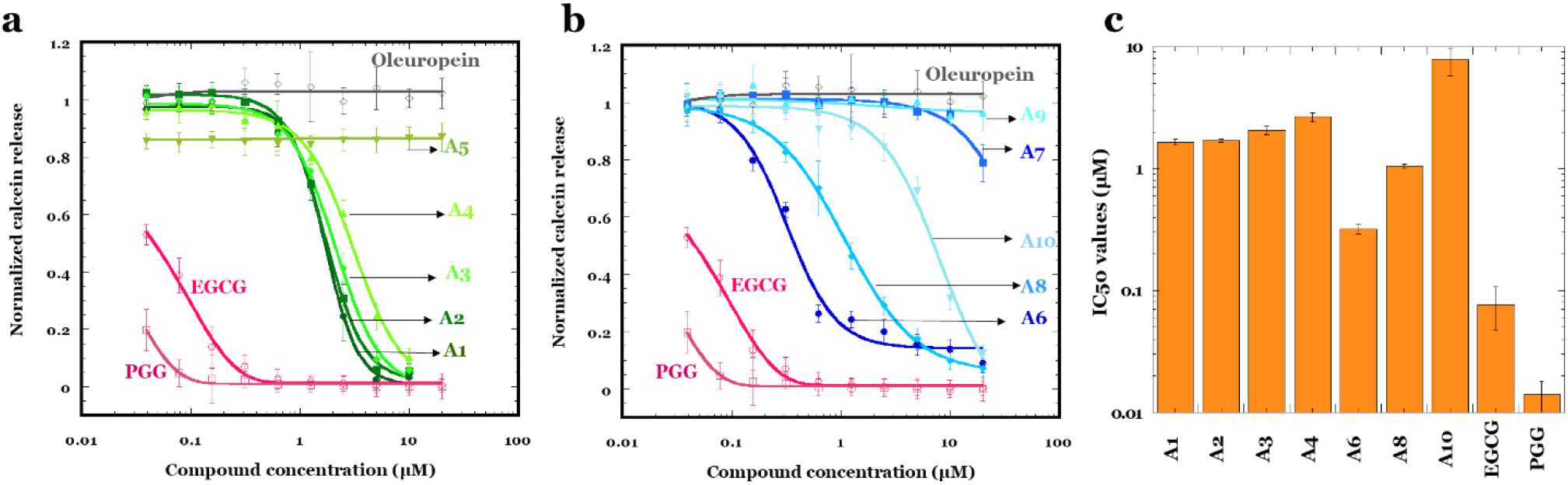
Quantifying the potency of lead compounds to inhibit membrane permeabilization. Dose-response calcein release for αSOs in presence of each of the hit compounds. All the values were normalized against calcein release due to αSOs alone in the absence of compounds. IC_50_ values were obtained by fitting dose-response curves to eq. 2. Note the logarithmic x-axes in panels a and b and the logarithmic y-axis in panel c.

### Using FIDA to elucidate the mechanism of binding inhibition

To investigate the underlying mechanism by which the compounds inhibit membrane permeabilization, we performed FIDA on αSOs – DOPG liposome mixtures in the presence of the hit compounds. For these experiments, 150 nM αSOs-Alexa488 and 5 µM compounds were pre-incubated for 15 mins, and 50 µM DOPG liposomes were then added after which the R_h_ was measured. (**Fig. 8**). Using the pre-determined αSO size (10.2 nm) as one of the species during taylorgram data fitting, we computed the percentages of liposome-bound and unbound species of oligomers based on the measured R_h_ values.

**Fig. 8.**
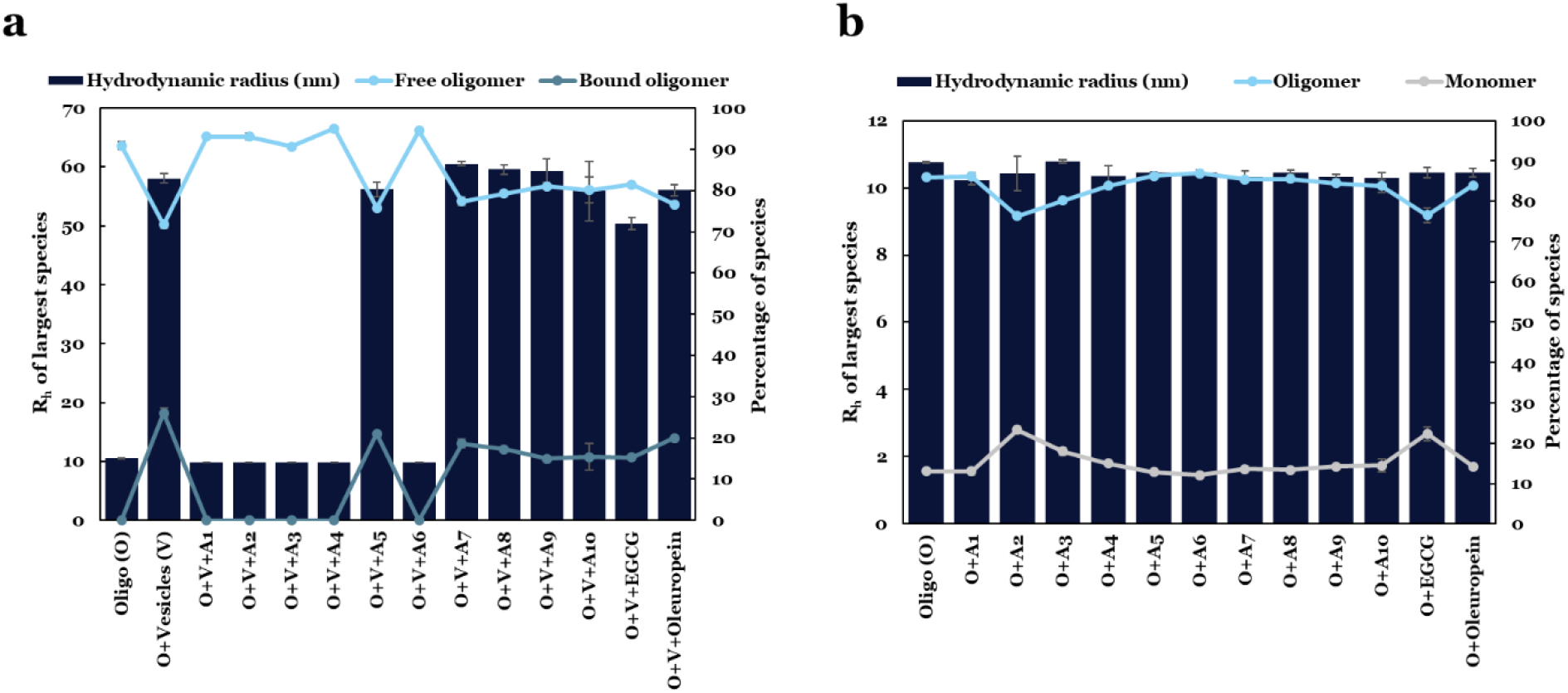
Interaction of αSOs with compounds in the presence and absence of liposomes analyzed by FIDA. a.) The apparent size (*R*_*h*_) of the largest species of αSOs-Alexa488 in the presence of DOPG liposomes and hit compounds. The values of *R*_*h*_ and percentage of species can be read on the left and right y-axes respectively. b.) *R*_*h*_ of free αSOs-Alexa488 (*i*.*e*. without liposomes) in the presence of the hit compounds.

The apparent size computed for αSOs-Alexa488 bound to liposomes was ∼60 nm. Remarkably, compounds A1, A2, A3, A4 and A6 completely prevented the formation of αSO:liposome complexes and retained the original size of αSOs (∼ 10 nm), indicating 0% bound oligomers (**Fig. 8a**). The remaining 5 compounds (A5, A7, A8, A9 and A10) did not completely prevent αSO – DOPG complex formation but showed a reduction in the percentage of bound species by 15-20% compared to control samples in the absence of hit compounds (26%).

While EGCG is more potent than all hit compounds in inhibiting calcein release during membrane permeabilization assay (IC_50_ = 0.08 µM), we note that the EGCG (unlike the 5 compounds A1-A4 and A6) could not completely displace αSOs from binding to DOPG liposomes, leading to ∼ 15% bound species (consistent with our previous observations^41^). The apparent size of liposome-bound αSOs in presence of EGCG was also reduced to 50 nm, which is slightly smaller than the apparent size of αSO-vesicle complex (∼60 nm) computed in the absence of hit compounds.

It is possible that inhibition of membrane permeabilization could also be a side effect due to complete dissociation of oligomers in to monomers, or aggregation of small soluble oligomers into less toxic larger oligomers in presence of hit compounds. We investigated this possibility in a new FIDA assay, where we incubated αSOs in presence of all 10 hit compounds along with two controls separately, and measured the sizes of labelled αSOs-Alexa488 (**Fig. 8b**). None of the compounds changed the size of the oligomer, demonstrating that the αSOs are intact and the inhibitory effect of compounds was not due to the dissociation or aggregation of αSOs. However, in presence of compounds A2, A3 and EGCG, the percentage of α-syn monomers were increased from 13% to 23%, 18% and 22% respectively (**Fig. 8b**).

To investigate the compounds’ own interactions with membranes, we incubated 100 µM of each compounds (A1 – A4, A6, A8 and A10) with and without 1 mM of DOPG vesicles for 30 minutes, ultracentrifuged the samples (1 h at 40,000 rpm), and subsequently determined the percentage of compound that is bound to vesicles using eq. 4. The five compounds A1, A2, A3, A4 and A6 that inhibited membrane permeabilization in the calcein assay, and also prevented the formation of αSO:liposome complexes in FIDA analysis, showed significant binding affinity to membrane with ∼60-95% of the compound pelleted along with vesicles (**Fig. 9**). On the other hand, compounds A8, A10 and EGCG that did not prevent the αSO – DOPG complex formation in FIDA assay but still inhibited membrane permeabilization in calcein assay showed no significant affinity to membrane, giving only ∼0-2% of bound compound in the pellet.

**Fig. 9.**
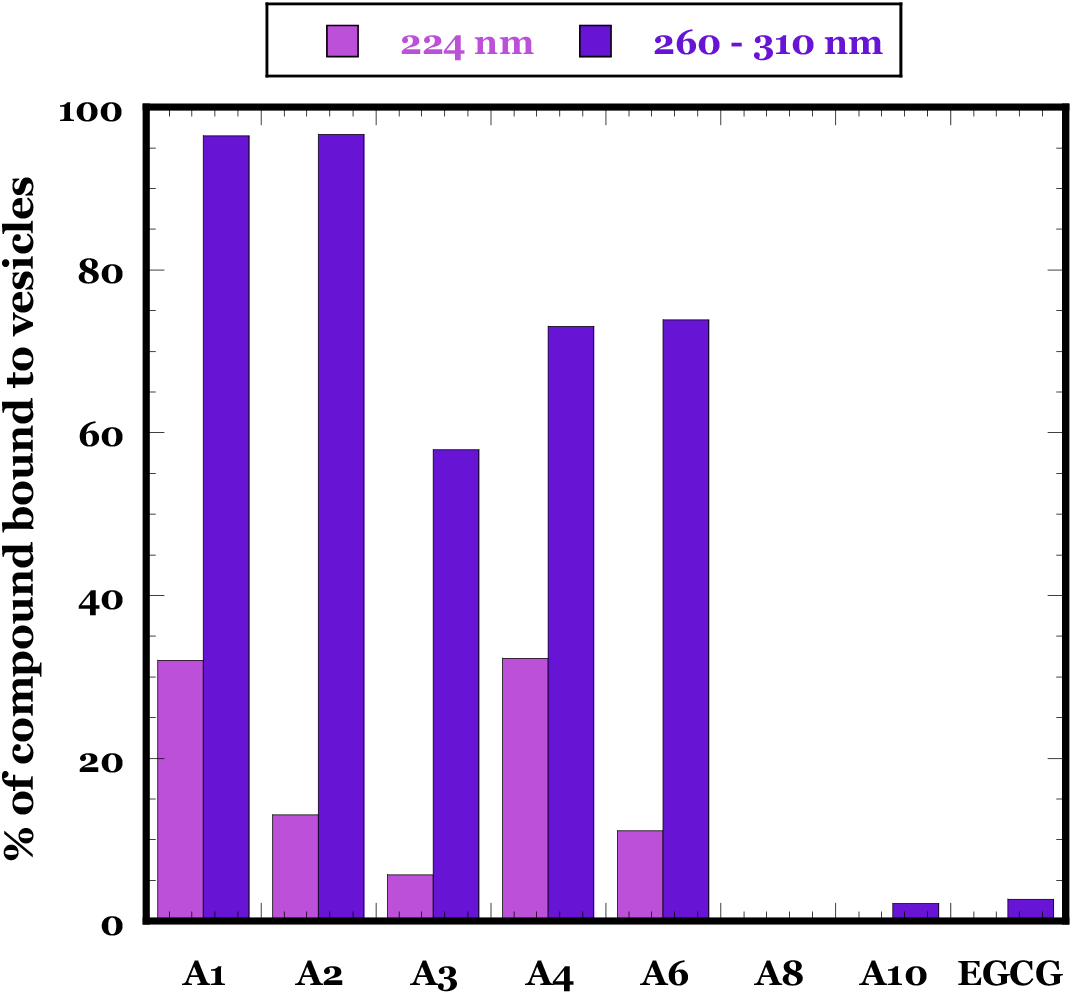
Percentage of compounds bound to DOPG vesicles. Data are based on change in absorption of 100 µM compounds in presence and absence of 1mM DOPG vesicles measured after incubating for 30 min and ultracentrifugation for 1 h at 40,000 rpm (eq. 4). Purple and blue refers to use of absorption at 224 nm and 260 – 310 nm respectively.

### The lead compounds show varying abilities to inhibit fibril formation

We finally evaluated whether the compounds identified from primary screening affect α-syn aggregation in ThT assays using 20 µM each of α-syn and compound. Among 10 lead compounds, compounds A5, A7 – A10 completely inhibited fibril formation similar to EGCG control at 20 µM. The 5 compounds that completely inhibited αSO binding to membranes (A1, A2, A3, A4 and A6) also altered the course of α-syn aggregation. Compounds A1, A2 and A3 decreased the half-time of fibrillation ∼2-fold compared to α-syn alone, while A4 and A6 modestly increased the lag phase (**Fig. 10**).

**Fig. 10.**
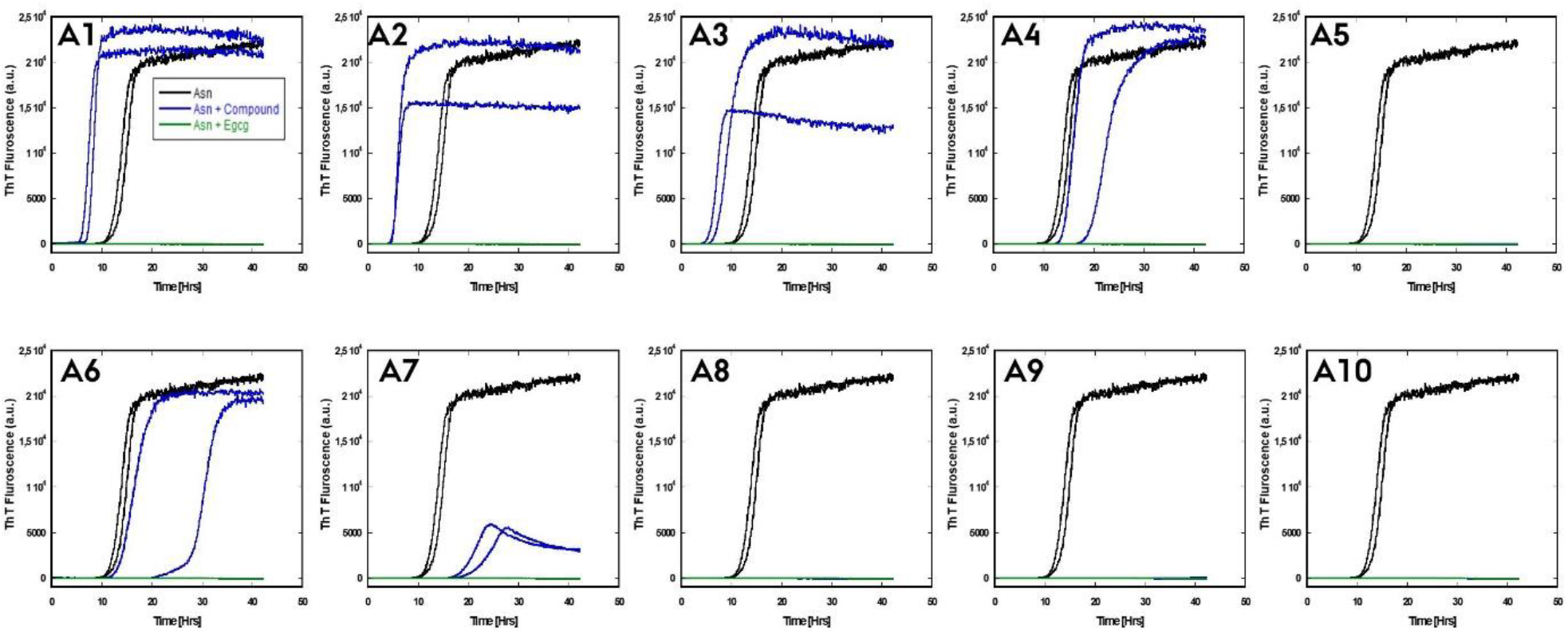
Efficiency of lead compounds in inhibiting fibrillation of 20 µM α-syn at equimolar ratio of compounds at 300 rpm shaking and 37 °C, monitored by ThT fluorescence.

### Sandwich ELISA identifies compounds that block αSO interactions with other proteins

To further demonstrate the utility of these hit compounds, sandwich ELISA of αSOs was performed. In this assay, nanobody 1 (NB1) (which is specific for *α*SO and does not bind monomer) was first immobilized on the plate which was then blocked, after which 1 µg/ml *α*SOs was added with or without 2 mM of each of the 10 hit compounds (A1 – A10). The concentration of αSO remaining bound after washing (which reflects the ability of the compound to inhibit αSO binding to either NB1, the antibody or both) was measured by adding an antibody raised against αSOs (14-9E7-A1). **Fig. 11a** summarizes the results. The positive control (absence of compounds) shows significant binding of αSOs to immobilized NB1, but several compounds (particularly A1 and A2, also A3 and A6 to a smaller extent) show a decrease in binding which indicate inhibition of oligomer contacts to either nanobody or antibody (or both). Compounds A4, A5, A7, and A10 lead to a slightly higher absorption than the positive control, due to background absorption of the compounds themselves upon binding. To investigate the best hits in more detail, we carried out a dose-response curve with A1 and A2 using NB1 and the monoclonal antibody 14-9E7-A1 (**Fig. 11b**). In both cases, there is a decline in absorption with increasing compound concentration as would be expected from a competition experiment, where the compound competes with NB1 or antibody for binding to *α*SOs. NB1 leads to a less scattered signal as well as a larger signal change, due both to a higher binding of *α*SOs in the absence of compound and (for A1) a lower level of binding at high compound concentrations. Fitting the NB1 data using an inverted binding isotherm leads to an apparent K_d_ of 0.13±0.04 and 0.12±0.09 mM for A1 and A2, respectively.

**Fig. 11.**
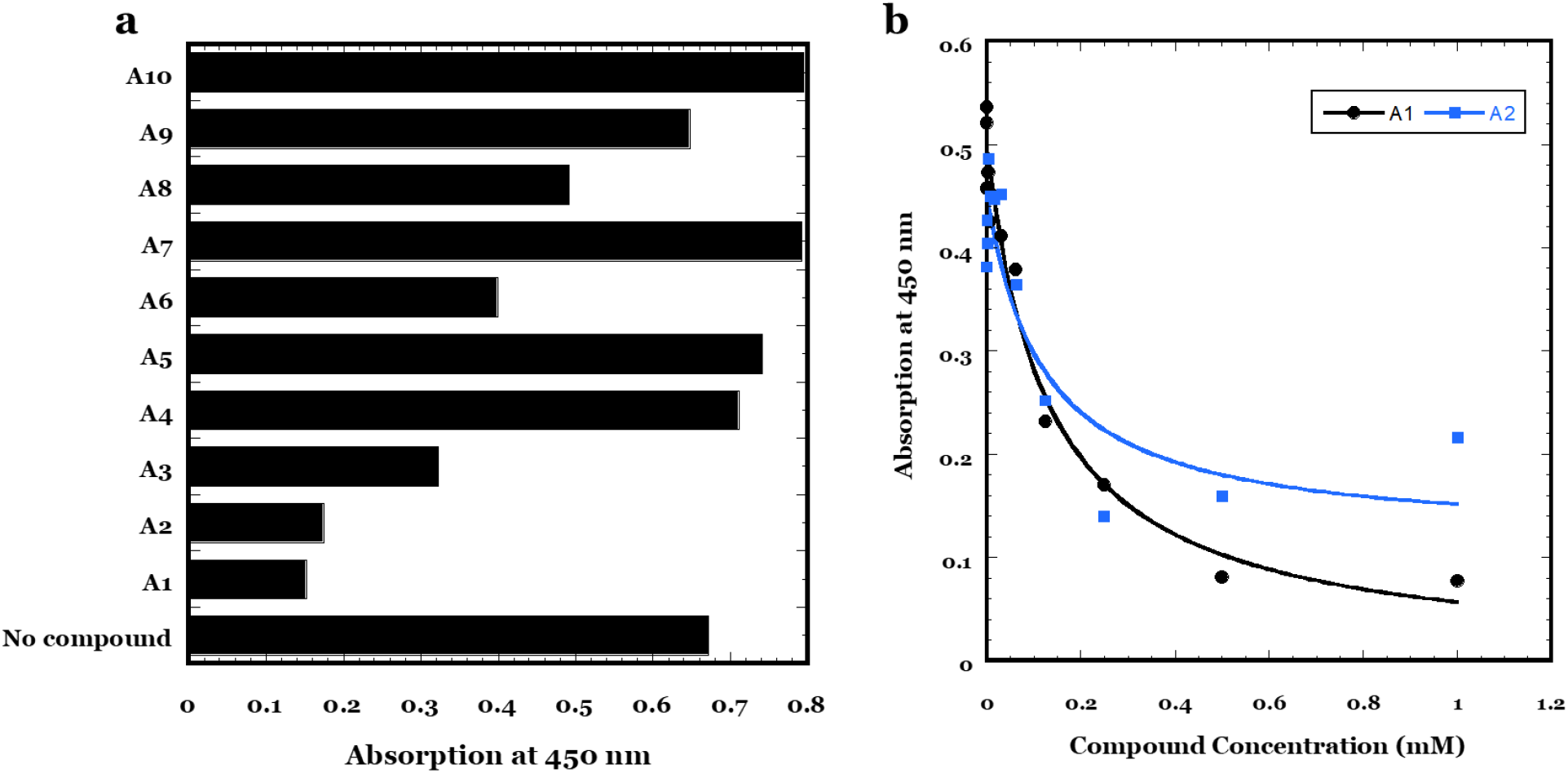
Sandwich ELISA to measure displacement of *α*SO from binding to nanobody NB1 and 14-9E7-A1 antibody. (a) Screening of all 10 hit compounds using 2 mM compounds and 1 µg/ml *α*SOs. (b) and (c) show dose response curves with 0-1 mM A1 and A2, respectively. Data are fitted to an inverted binding isotherm to give an apparent K_d_ of 0.13±0.04 and 0.12±0.09 mM for A1 and A2, respectively.

## DISCUSSION

Existing treatments of PD mainly target dopamine-related symptoms by either increasing the amount of dopamine using dopamine agonists or inhibiting endogenous enzymes that break down dopamine. Therefore, drugs that can reduce neuronal toxicity and increase neuronal viability are vital in treating PD. Since αSOs are more pathogenic on a per-mass basis than α-syn monomers and fibrils, identification of small molecules that can target oligomer-derived cytotoxicity might eventually reduce pathogenesis of PD. Here, we established a simple assay to screen small molecules that prevent interactions of αSOs with cell membranes as a proxy for the toxic mechanisms of αSOs in the cell.

### Protein ligands bind transiently to αSOs but significantly promote monomer aggregation

We first identified a small collection of potential αSO binding ligands based on previous proteomics studies. We then investigated the strength of αSOs interactions with these proteins using different biophysical techniques to evaluate their potential use in a small molecule screening assay. It is evident from both dot blot and SPR studies (where α-syn species were applied at concentrations of 7 µM, *i*.*e*. well below the 150 µM used in solution studies) that these protein ligands exhibits strong binding interaction when one of the binding partner is immobilized on a surface. In contrast, the affinity of ligands to αSOs in solution is so weak as not to be detected when measured by FIDA, Note that all the ligands that are identified using co-immunoprecipitation have αSOs immobilized on agarose beads using α-syn-specific antibodies. This suggests in line with other reports that immobilization can lead to artifactual results, both false negatives^43^ and false positives^44^ caused *e*.*g*. by surface-assisted avidity effects^45^ or hydrophobic contacts to the surface^46^. In addition, structural differences and changes in conformational flexibility of the protein ligands when they are in solution and bound to a surface may also determine the binding of αSOs.

In summary, our FIDA data suggest the interactions between the oligomer and the four ligands (Cfl1, Uchl1, Sirt2 and SerRS) are very weak when both binding partners are in solution. In addition, it is evident from dot-blot densitometric analysis that α-syn monomer shows similar or weaker binding to all four ligands when compared to αSOs. It is noteworthy that three of the four protein ligands dramatically increased α-syn aggregation during shaking-induced aggregation assays. It is well known that weakly transient bound protein complexes play a crucial role in cell metabolism as well as in regulatory and signaling pathways with affinities in the µM-mM range ^47,48^. The implication is that the protein ligands do not form stable complexes with αSOs but change the course of aggregation through transient interactions with other α-syn species, most likely the monomer.

### Investigating the pathological interactions of αSOs and cell membranes

The physiological role of α-syn-membrane interactions is strongly associated with the ability of α-syn to cluster synaptic vesicles and chaperones SNARE complex formation to maintain neurotransmitter release^49^. These interactions are primarily driven by the lipophilic initial 25 residues in the N-terminal region, where the first 14 residues penetrates into lipid head groups as anchor, leaving the remaining residues folded as an α-helix on the membrane surface^7,50^. Such αSOs-membrane interactions can turn pathological in the case of the αSO, where N-terminal binding is accompanied by insertion of a rigid β-sheet rich oligomer core into the lipid membrane, thereby disrupting its integrity. This further triggers a channel like pore formation and disrupts cellular calcium ion homeostasis, leading to cell death^5,8^.

FIDA and the calcein assay allowed us to quantify binding affinity between αSOs-Alexa488 and DOPG vesicles, resulting in an apparent 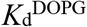 of ∼40 µM and 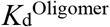 of ∼143 nM respectively. The apparent discrepancy reflects both the different types of assays (direct in FIDA and indirect in the calcein assay) and the fact that 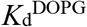 is based on monomeric lipid units; it would probably be more appropriate to express lipid concentration in terms of available binding sites but that is not possible to gauge. However, the ratio between the two *K*^d^ values (ca. 280) would at face value suggest a ratio of 280 lipid molecules per oligomer which is not an unreasonable figure, given that 280 DOPG molecules have a molecular weight of 217 kDa which is in the same ball-park region as the αSO (450 kDa).

Different conformations of α-formed during the course of α-syn aggregation have different binding affinity towards cell membranes. For example, the later-stage α-syn oligomers (“type B”) identified by Fusco *et al* showed the greatest membrane affinity, followed by α-syn monomers, early-stage (“type A”) oligomers and finally fibrils^8^. Type A and B oligomers have similar sizes and morphologies but different abilities to disrupt lipid bilayers, clearly linked to their different structural features. We exploited this disruptive αSO:membrane interaction to establish a screening assay, and screened two datasets of small molecules.

### General considerations about hit compounds

Both compound sets used in our primary screening are quite diverse, and the promising hit compounds are found in different structural classes. Nevertheless some groupings are possible.

Compounds A1 – A3 and A6 share the central feature of having two aromatic rings connected by an alkyl chain. These compounds show a broad range of activity in their original application, including antimicrobial and surfactant properties (chlorhexidine, methyl benzethonium, benzethonium, metacycline and shikonin), antispasmodics that block calcium channels (pinaverium), antagonists of acetylcholine receptors (methoctramine), chemotherapy medication which binds DNA and stops replication (cisplatin and carboplatin) and a compound produced by the oxidation of adrenaline which affects mood and thought processes (adrenochrome). Though none of these compounds have been studied directly in relation to αSOs before, some of them have been reported with reference to other aspects of neurodegeneration. For example, methoctramine competitively antagonizes acetylcholine receptors (M2 muscarinic receptors), thus improving memory in cognitively impaired aged rats^51^. Chlorhexidine is an inhibitor of the Keap1 regulator, thus boosting the antioxidant potential of dopaminergic neurons^52^. Cisplatin, a Pt^II^ containing antitumor drug, has been shown to inhibit α-syn aggregation by coordinating platinum to side chains of methionine and histidine residues^53^. In addition, metacycline is an antibiotic, shikonin a quinone and adrenochrome has also been shown to reduce α-syn aggregation^54,55^.

### Hit compounds and their various courses of action in preventing membrane disruption of αSOs

Out of 11 hit compounds identified in the primary screen, we confirmed 7 compounds (ranked by impact as A6>A8>A1=A2=A3=A4>A10) in a more detailed dose-response study, while A5, A9 and A10 failed to show any effect. A1, A2, A3, A4 and A6 all inhibited αSOs binding to membranes to an equal extent (whereas the remaining 5 compounds had no effect compared to the oligomer alone) in an orthogonal FIDA assay based on oligomer size. These 5 effective compounds all showed significant binding affinity towards membranes on their own (ranked A1=A2>A4=A6>A3). This suggests that these compounds compete with oligomers for membrane binding, enabling them to either displace them from, or (equivalently) prevent them from binding to, the surface of membranes. The 5 compounds also block αSO contacts with anti/nanobodies in our ELISA sandwich assay. This implies that they inhibit interactions either at the oligomer-antibody or oligomer-nanobody interface (or both). Together with their membrane-binding and leakage-blocking activity, thus highlights their versatility, *i*.*e*. multiple mechanisms of αSO containment. The membrane blocking phenomenon has previously been observed for several aminosterol compounds such as squalamine and trodusquemine, which in this way are thought to suppress the toxicity of Aβ and αSOs^19,56,57^. Claramine, a blood–brain barrier permeable small molecule from the aminosterol class, has also shown to prevent the toxicity of pore forming agents including melittin from honeybee venom and α-hemolysin from *Staphylococcus aureus*^*58*^. Compounds A8 and A10 reduced the amount of calcein release but did not affect binding to DOPG liposomes, suggesting that the compounds could perhaps reduce oligomer membrane permeabilization without preventing binding (*e*.*g*. by leading to more superficial membrane attachment), as observed for the control compound EGCG^41^. However, with both compounds the oligomers retain the same overall size, indicating that they do not cause either dissociation or aggregation of oligomers (**Fig. 12**).

**Fig. 12.**
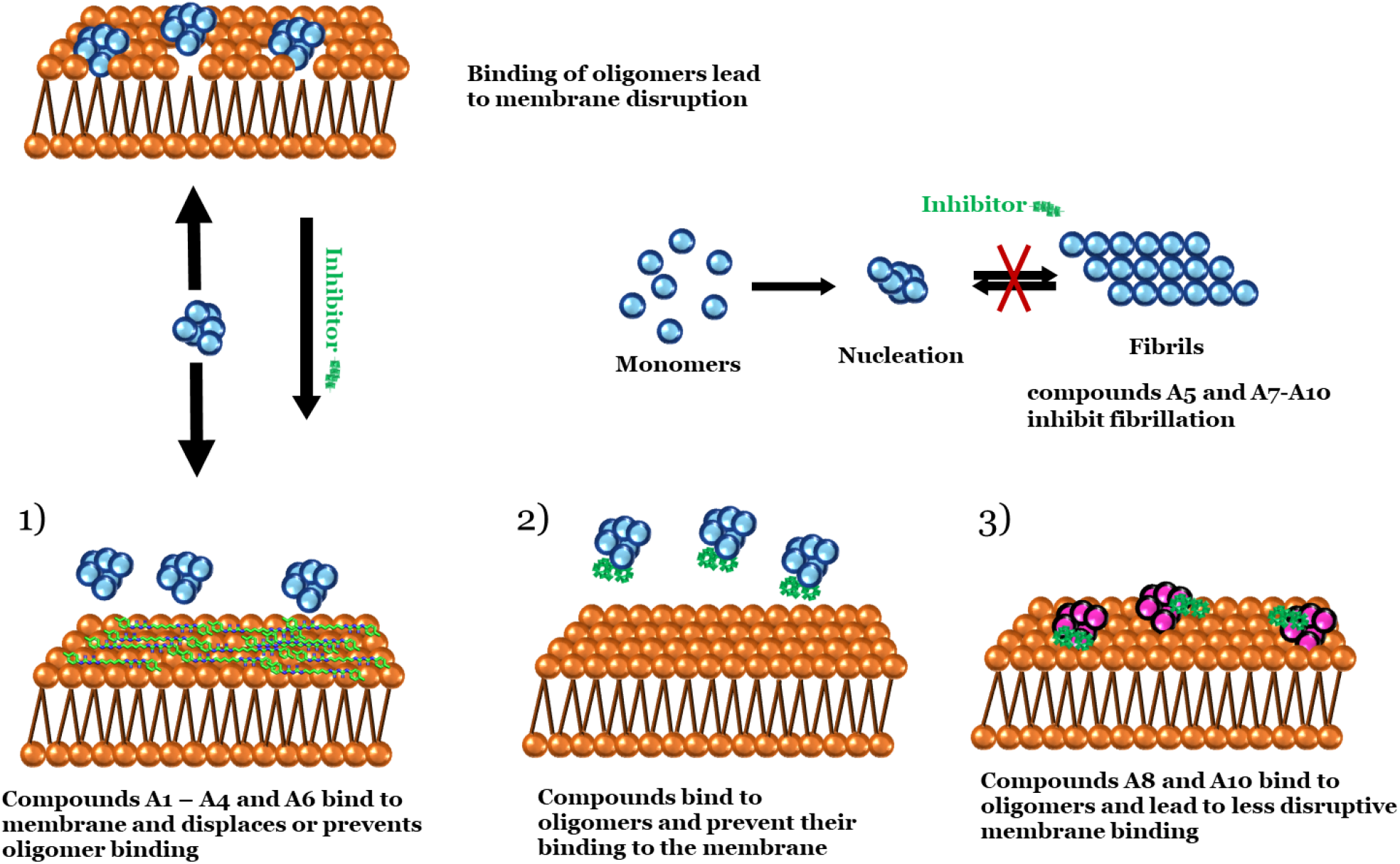
Schematic representation of hit compounds inhibiting membrane permeabilization and fibrillation.

Finally, it is noteworthy that the roles are completely reversed in our fibrillation assays: here compounds showing poor performance against αSO (A5 and compounds A7-A10) are very effective at inhibiting fibrillation while the remaining 5 compounds show only a modest increase in the lag phase (A4 and A6) or even an acceleration of fibrillation (A1-A3). This complementarity of action suggests that different mechanisms are required to target the αSO and the α-syn fibril, again implying that the two targets have substantially different structural properties and thus different binding interfaces. Such molecular insight may be useful in guiding future therapeutic strategies.

## Abbreviations

*E. coli*: *Escherichia coli*
α-syn: α-synuclein monomer
αSO: α-synuclein oligomer
ThT: Thioflavin T
SPR: Surface Plasmon Resonance
FIDA: Flow Induced Dispersion Analysis

## Acknowledgements

A.K.S. is supported by the Lundbeck Foundation (grant R287-2018-1836). D.E.O. is supported by the Independent Danish Research Council | Natural Sciences (grant 8021-00208B, funding M.N.), Novo Nordisk Foundation (grant NNF17OC0028806, funding P.A.) and the Lundbeck Foundation (grant R276-2018-671).

